# Mass spectrometric profiling of estrogen and estrogen metabolites in human stool and plasma partially elucidates the role of the gut microbiome in estrogen recycling

**DOI:** 10.1101/2024.08.07.606543

**Authors:** Vince W. Li, Tien S. Dong, Diana Funes, Laura Hernandez, Srinivasa T. Reddy, Emeran Mayer, Lin Chang, David Meriwether

## Abstract

Estrogen and estrogen metabolites are commonly measured in human plasma and serum, but there exist almost no reports of estrogen measured in human stool. This methodological limitation in turn limits our understanding of the relationship between systemic and intestinal estrogen. We thus developed a highly sensitive liquid chromatography-mass spectrometry/mass spectrometry (LC-MS/MS) method for measuring free and conjugated forms of 15 estrogens and estrogen metabolites in human stool and plasma. We first investigated human stool and plasma estrogen in healthy control males; follicular and luteal phase premenopausal females; and postmenopausal females. Most estrogens were present in the plasma and stool of all groups, and plasma estrogen levels correlated with stool estrogen levels. In stool, estrogens were higher in premenopausal females, with estrogen levels rising across the menstrual cycle. We further combined these measures with shotgun metagenomic sequencing of the stool microbiomes. The level of estrogen deconjugation enzyme gene copy number (beta-glucuronidase + arylsulfatase) was higher in premenopausal females; while the gene copy numbers of beta-glucuronidase + arylsulfatase, but not beta-glucuronidase alone, correlated with reactivated stool estrogen in all groups. Moreover, deconjugation enzyme gene copy number correlated with plasma total estrogen in males and with individual plasma estrogen metabolites in all groups. These results support the hypothesis that gut microbial beta-glucuronidase and arylsulfatase control the reactivation of gut estrogen while modulating systemic levels through the uptake and recirculation of reactivated estrogen.

## 1. Introduction

The primary estrogens estradiol and estrone can be modified during phase I metabolism by hydroxylation of carbons 2, 4, and 16, while the hydroxyls at carbons 2 and 4 can be further methylated during phase II metabolism. Analytical LC-MS/MS methods for the determination of primary estrogen and estrogen metabolites have been validated for human serum, plasma, and urine; and these methods have shed light on important aspects of the biology of estrogen metabolites. However, during phase II metabolism estrogens are also conjugated to sulfate and glucuronide moieties; and these estrogen conjugates are excreted not only into urine but also into the gut lumen. Surprisingly, there exist no published validated LC-MS/MS methods for the determination of primary estrogens and estrogen metabolites in human stool; and there exist no published datasets that combine estrogen and estrogen metabolite data from human stool with data from human plasma or serum. This limitation of prior research introduces important knowledge gaps concerning the relationships between circulating estrogens, intestinal estrogens, and gut microbiota. We have developed and validated an LC-MS/MS method for determining primary estrogen and estrogen metabolite levels in human stool and plasma. We analyze the stool and plasma of healthy control males and pre- and postmenopausal females, and we combine this estrogen data with stool shotgun metagenomic sequencing to address some of the gaps in knowledge we have identified.

### 1.1. Estrogens and estrogen biology

Estrogens are sex steroid hormones that play an essential role in the development and function of the female reproductive system [1, 2], but they influence various additional aspects of health and well-being in both sexes [3, 4]. Estrogens can affect mood [5], metabolism [6], bone density [7], cardiovascular health [8], and cognitive functions [9]; and there are important links between estrogens and estrogen metabolites and various cancers [10] including ovarian cancer [11, 12], breast cancer [13], and even prostate cancer [14].

Estrone (E1), estradiol (E2), and estriol (E3) are the primary bioactive estrogens. Estradiol has the highest binding affinity for the estrogen receptors ERα and Erβ, while estrone and estriol exhibit relative binding affinities of approximately 50% and 15% in comparison [15]. Estradiol is the dominant estrogen in premenopausal women, and it is synthesized in the ovaries by aromatization of androstenedione followed by dehydrogenation of estrone (**Figure 1A**) [16]. Estrone is the primary estrogen in males and postmenopausal women, and both estrone and estradiol can be synthesized from testosterone and androstenedione in non-ovarian tissues like adrenal glands and adipose tissue [17]. By contrast, estriol—important during pregnancy—is formed in the placenta following phase I metabolism of E1 and E2 by cytochrome P_450_ (CYP450)-mediated oxidation of carbon 16 in the steroid ring (**Figure 1A**). CYP450 enzymes present in breast and other estrogen-target tissue can also hydroxylate E1 and E2 at carbons 2 and 4, producing 2- and 4-OH-metabolites (**Figure 1A**) [13]. These 2- and 4-OH-metabolites—collectively termed *catechol estrogens*—have lower affinity for the estrogen receptors than do their primary estrogens, and thus hydroxylation is a first step in the estrogen degradation and excretory pathway [18]. However, catechol estrogens can be further oxidized to quinones and semi-quinones that can form reactive oxygen species that damage DNA and lipids [19]; and higher levels of 4-OHE1 have been associated with increased risk of breast cancer [20] and breast cancer invasiveness [21].

**Figure 1.**
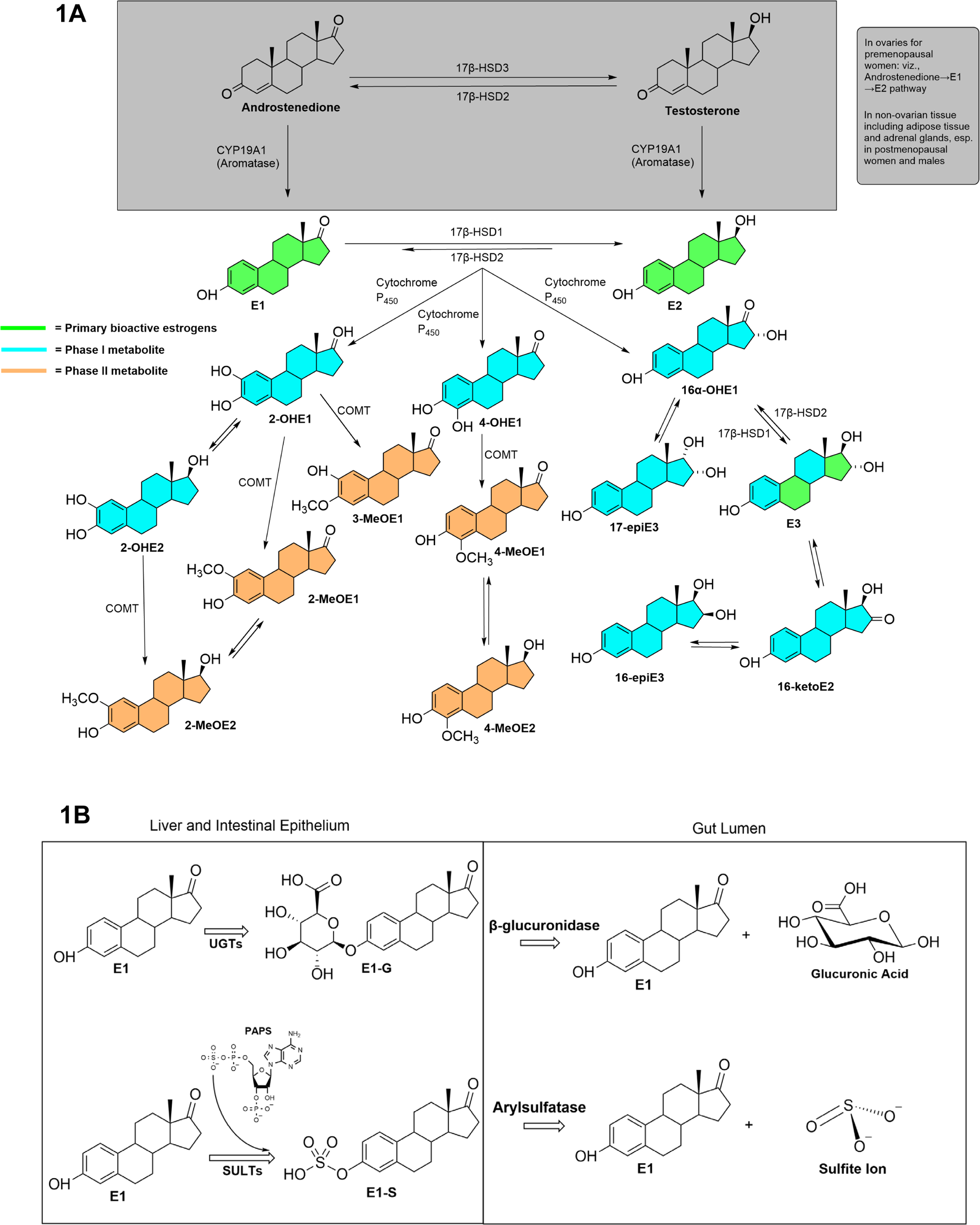
Synthesis and gut microbial catabolism of phase I and phase II metabolites of estrogen. **(A)** Synthesis pathway of estrone and estradiol together with metabolic pathways of their phase I hydroxylation and phase II methoxylation. (**B**) Phase II sulfation and glucuronidation of estrogens (*left panel*) together with catabolism of estrogen conjugates by gut microbial β-glucuronidase and arylsulfatase enzymes *(right panel*). Figures 1A and 1B adapted from [1–5]. **<u>Abbreviations</u>**: **16-epiE3**, 16-epiestriol; **16-ketoE2**, 16-ketoestradiol; **16α-OHE1**, 16α-hydroxyestrone; **17β-HSD**, 17β-Hydroxysteroid dehydrogenase; **17-epiE3**, 17-epiestriol; **2-MeOE1**, 2-methoxyestrone; **2-MeOE2**, 2-methoxyestradiol; **2-OHE1**, 2-hydroxyestrone; **2-OHE2**, 2-hydroxyestradiol; **3-MeOE1**, 3-methoxyestrone; **4-MeOE1**, 4-methoxyestrone; **4-MeOE2**, 4-methoxyestradiol; **4-OHE1**, 4-hydroxyestrone; **COMT**, catechol-o-methyltransferase; **CYP19A1**, cytochrome P450 family 19 subfamily A member 1; **CYP450,** cytochrome P_450_; **E1**, estrone; E1-G, estrone glucuronide; E1-S, estrone sulfate; **E2**, estradiol; **E3**, estriol; **PAPS**, 3’-phosphoadenosine-5’-phosphosulfate; **UGT**, uridine 5’-diphospho-glucuronosyltransferase; **SULT**, sulfotransferases.

The catechol estrogens are further inactivated by the phase II metabolic enzyme catechol O-methyl transferase (COMT), which catalyzes methylation at carbons 2, 3, or 4 (**Figure 1A**) [18]. The resultant methoxyestrogens lack further oxidative and genotoxic potential and possess little to no affinity for ERα and ERβ [22]. However, methoxyestrogens resist additional catabolism and like other estrogen species are lipophilic, poorly water soluble, and difficult to excrete. Thus, further phase II metabolism of estrogen produces sulfate and glucuronide conjugates (**Figure 1B**). Estrogen sulfotransferase (SULT) mediates the transfer of a sulfate moiety from the cofactor 3’-phosphoadenosine-5’-phosphosulfate (PAPS) to the carbon 3 or 17 via reaction with 3-OH or 17-OH [18]. UDP glucuronosyltransferase (UGT) present in the liver, kidney, and intestine likewise catalyzes the conjugation of estrogens with glucuronic acid at a hydroxyl to form estrogen 3-, 16-, or 17-glucuronides [23]. The increased hydrosolubility of the estrogen sulfates and glucuronides prevents their binding to estrogen receptors and makes them more suitable for excretion [24]. Estrogen sulfates and glucuronides are subsequently excreted into urine, into bile, or transintestinally directly into the gut lumen via ATP binding cassette (ABC) transporters located in the kidney, liver hepatocytes, and intestinal enterocytes [25].

Human gut microbiota produce β-glucuronidase [26] and arylsulfatase enzymes [27], and it has been demonstrated in vitro that gut microbial β-glucuronidase can metabolize estrogen glucuronides and reactivate estrogen [28]. It has thus been hypothesized that gut microbial reactivation of estrogen (**Figure 1B**) can modulate systemic estrogen levels via enterohepatic recirculation or recycling [29, 30].

### 1.2. Determination of estrogen levels in human serum, plasma, and urine

Given the importance of estrogens for human health and disease, various analytical methods have been developed to determine the levels of estrogens in human serum, plasma, and urine (reviewed in [31]). Enzyme linked immunoassays (ELISAs) remain in common use in clinical labs due to their low cost and routine nature, but they fail to overcome several technical challenges associated with estrogen measurement in males and postmenopausal females, and with measurement of estrogen metabolites in addition to primary estrogens. First, the circulating concentrations of estrogens especially in males and postmenopausal females are low: the reference range of e.g. estradiol is only 30 to 400 pg/mL for premenopausal women, but an even lower 0 to 30 and 10 to 50 pg/mL for postmenopausal women and men [32, 33]. A successful analytical method must therefore be highly sensitive. Second, many estrogen species especially metabolites overlap in both molecular weight and chemical structure (e.g., 2-OHE1 and 4-OHE1). A successful analytical method must therefore also be highly specific. However, antibody-based detection methods lack specificity at low concentration; are unable to accurately measure estrogen levels in men and postmenopausal women; and are commonly limited to estrone and estradiol while failing to resolve estrogen metabolites [31].

Liquid chromatography-mass spectrometry/mass spectrometry (LC-MS/MS) combined with chemical derivatization during sample preparation can successfully overcome these challenges of sensitivity and specificity. While estrogens themselves ionize poorly [34], estrogens that have been chemically modified through reaction with derivatizing agents like dansyl chloride (DSC) or pyridine-3-sulfonyl chloride can yield limits of detection and quantitation of approximately 1 pg/mL [31]. Moreover, the addition of highly resolving small particle liquid chromatography with MS/MS can overcome the challenge of isobaric overlap between estrogen species and enable the highly specific determination of not just primary estrogens but of estrogen metabolites as well.

LC-MS/MS based determinations of primary estrogen and estrogen metabolites in plasma and serum of pre- and postmenopausal women have produced important insights about systemic estrogen metabolites and overall health and disease. For example, postmenopausal women with increased 2-hydroxylation of estrone exhibited lower body fat and increased lean body mass [35]; while postmenopausal women with increased 4-hydroxylation of estrone were at greater risk of breast cancer and breast cancer invasiveness [20, 21]. In brief, LC-MS/MS based determinations of systemic estrogens have shed light on important aspects of estrogen biology.

### 1.3. Limitations of prior research and gaps in knowledge

To our knowledge, there exists only a single report of the LC-MS/MS determination of primary estrogens and estrogen metabolites in human stool [36]. However, the authors adapted an LC-MS/MS method that had been validated for human serum [37] and urine [38] without explicitly validating the method for stool; and they combined determination of estrogen in stool with determination of estrogen in urine but not plasma or serum. Otherwise, we are aware of no further reports of LC-MS/MS determination of primary estrogens and estrogen metabolites in human stool, and no presentation of stool estrogen together with circulating estrogen data.

This limitation of prior research generates several important gaps in knowledge regarding the relationships between systemic estrogens, intestinal estrogens, and gut microbiota.

1. Estrogen sulfates and estrogen glucuronides can be excreted into the gut via ABC transporters located in hepatocytes and enterocytes [25]. However, the mechanism by which the excretion rate of conjugated estrogens is determined is not clear. Is excretion rate into the gut a simple function of circulating estrogen concentration, or is there an additional circuit that determines the rate of estrogen excretion? Positive correlation between systemic estrogen level and stool estrogen level would support the first hypothesis.
2. Human gut microbiota can produce both β-glucuronidase and arylsulfatase enzymes, but research into the reactivation of gut estrogen has largely focused on β-glucuronidase [27, 36]. Is gut microbial β-glucuronidase largely responsible for reactivation of gut estrogen or do arylsulfatase enzymes also play a role? Determining correlations between reactivated estrogen in stool and β-glucuronidase versus β-glucuronidase + arylsulfatase would shed light on this question.
3. It is commonly assumed that gut reactivated estrogens can be reabsorbed enterohepaticaly and can thereby elevate circulating estrogen levels [28–30]. However, direct evidence for this hypothesis is weak. Flores et al. showed that there exists a positive correlation between gut microbial beta glucuronidase activity and urinary estrone in men [36]. However, urinary estrogen levels only weakly correlate with circulating estrogen levels [39]; and the contribution of arylsulfatase was not evaluated. Importantly, we know of no study that examined the relationships between gut microbial β-glucuronidase and arylsulfatase, human stool estrogen, and plasma or serum estrogen.

### 1.4. Project overview

We sought to overcome the limitations of prior research by first developing and validating an LC-MS/MS method for the determination of free and conjugated primary estrogens and estrogen metabolites in human stool and plasma. We adapted a method that had been previously validated in serum [37], but we employed in-house synthesized analyte-specific stable heavy isotope internal standards to overcome the matrix effects of stool and improve precision and accuracy. There exist no prior reports of an LC-MS/MS method validated for determining estrogen levels in stool, so we present the results of our development and validation studies here.

We next employed this method to determine the levels of free and conjugated estrogens in the plasma and stool of healthy control males, follicular and luteal premenopausal females, and postmenopausal females. Our plasma results are within the reference range for each group. We present novel data on sex and menstrual-specific differences in human stool estrogen level, while also providing for the first time data on the relationship between stool and plasma level.

Finally, we combined our estrogen data with determination of gut microbial β-glucuronidase and arylsulfatase gene copy numbers by shotgun metagenomic sequencing. We thereby provide evidence that both β-glucuronidase and arylsulfatase together are responsible for reactivation of stool estrogen, and that both enzymes together might modulate systemic estrogen levels via reactivation and recycling of stool estrogens.

## 2. Materials and Methods

### 2.1 Calibrants

Fifteen estrogens and estrogen metabolites including estrone (E1), estradiol (E2), estriol (E3), 2-hydroxyestrone (2-OHE1), 2-hydroxyestradiol (2-OHE2), 4-hydroxyestrone (4-OHE1), 16α-hydroxyestrone (16α-OHE1), 3-methoxyestrone (3-MeOE1), 4-methoxyestrone (4-MeOE1), 17-epiestriol (17-epiE3), 2-methoxyestrone (2-MeOE1), 2-methoxyestradiol (2-MeOE2), 4-methoxyestradiol (4-MeOE2), 16-epiestriol (16-epiE3), and 16-ketoestradiol (16-ketoE2) were purchased from Steraloids, Inc. (Newport, RI). Deuterium-labeled estrogen metabolites including estrone-2,4,16,16-d4 (E1-d4) and estradiol-2,4,16,16-d4 (E2-d4) were purchased from Steraloids, Inc (Newport, RI). See Table 1 for IUPAC names. All deuterated and non-deuterated estrogen analytical standards were reported ≥ 98% chemical and isotopic purity.

**Table 1.** Primary estrogen and estrogen metabolite targets: IUPAC name, common name, abbreviation; together with monoisotopic mass of each parent and derivatized estrogen.

| <u>IUPAC Name</u> | <u>Common Name</u> | <u>Abbreviation</u> | <u>Monoisotopic Mass (Mmi)</u> | <u>E-DS Mmi</u> | <u>E-DS-D6 Mmi</u> |
| --- | --- | --- | --- | --- | --- |
| <u>3-Hydroxyestra-1,3,5(10)-trien-17-one</u> | <u>Estrone</u> | <u>E1</u> | <u>270.16</u> | <u>503.19</u> | <u>509.19</u> |
| <u>3-Hydroxyestra-1,3,5(10)-trien-17-one-2,4,16,16-D4</u> | <u>Estrone-2,4,16,16-D4</u> | <u>E1-D4</u> | <u>274.16</u> | <u>507.19</u> | <u>513.19</u> |
| <u>Estra-1,3,5(10)-triene-3,17β-diol</u> | <u>Estradiol</u> | <u>E2</u> | <u>272.18</u> | <u>505.21</u> | <u>511.21</u> |
| <u>Estra-1,3,5(10)-triene-3,17β-diol-2,4,16,16-D4</u> | <u>17β-Estradiol-2,4,16,16-D4</u> | <u>E2-D4</u> | <u>276.18</u> | <u>509.21</u> | <u>515.21</u> |
| <u>Estra-1,3,5(10)-triene-3,16α,17β-triol</u> | <u>Estriol</u> | <u>E3</u> | <u>288.17</u> | <u>521.20</u> | <u>527.20</u> |
| <u>2,3-Dihydroxyestra-1,3,5(10)-trien-17-one</u> | <u>2-Hydroxyestrone</u> | <u>2-OHE1*</u> | <u>286.16</u> | <u>519.19</u> | <u>525.19</u> |
|  |  |  |  | <u>752.22*</u> | <u>764.22*</u> |
| <u>Estra-1,3,5(10)-triene-2,3,17β-triol</u> | <u>2-Hydroxyestradiol</u> | <u>2-OHE2*</u> | <u>288.17</u> | <u>521.20</u> | <u>527.20</u> |
|  |  |  |  | <u>754.23*</u> | <u>766.23*</u> |
| <u>3,4-Dihydroxyestra-1,3,5(10)-trien-17-one</u> | <u>4-Hydroxyestrone</u> | <u>4-OHE1*</u> | <u>286.16</u> | <u>519.19</u> | <u>525.19</u> |
|  |  |  |  | <u>752.22*</u> | <u>764.22*</u> |
| <u>3,16α-Dihydroxyestra-1,3,5(10)-trien-17-one</u> | <u>16α-Hydroxyestrone</u> | <u>16α-OHE1</u> | <u>286.16</u> | <u>519.19</u> | <u>525.19</u> |
| <u>3-Methoxyestra-1,3,5(10)-trien-17-one</u> | <u>3-Methoxyestrone</u> | <u>3-MeOE1</u> | <u>300.17</u> | <u>533.20</u> | <u>539.20</u> |
| <u>3-Hydroxy-4-methoxyestra-1,3,5(10)-trien-17-one</u> | <u>4-Methoxyestrone</u> | <u>4-MeOE1</u> | <u>300.17</u> | <u>533.20</u> | <u>539.20</u> |
| <u>Estra-1,3,5(10)-triene-3,16α,17α-triol</u> | <u>17-Epiestriol</u> | <u>17-epiE3</u> | <u>288.17</u> | <u>521.20</u> | <u>527.20</u> |
| <u>3-Hydroxy-2-methoxyestra-1,3,5(10)-trien-17-one</u> | <u>2-Methoxyestrone</u> | <u>2-MeOE1</u> | <u>300.17</u> | <u>533.20</u> | <u>539.20</u> |
| <u>1,3,5(10)-triene-3,17β-diol</u> | <u>2-Methoxyestradiol</u> | <u>2-MeOE2</u> | <u>302.19</u> | <u>535.22</u> | <u>541.22</u> |
| <u>4-Methoxyestra-1,3,5(10)-triene-3,17β-diol</u> | <u>4-Methoxyestradiol</u> | <u>4-MeOE2</u> | <u>302.19</u> | <u>535.22</u> | <u>541.22</u> |
| <u>1,3,5(10)-Estratriene-3,16β,17β-triol</u> | <u>16-Epiestriol</u> | <u>16-epiE3</u> | <u>288.17</u> | <u>521.20</u> | <u>527.20</u> |
| <u>3,17β-Dihydroxyestra-1,3,5(10)-trien-16-one</u> | <u>16-Ketoestradiol</u> | <u>16-ketoE2</u> | <u>286.16</u> | <u>519.19</u> | <u>525.19</u> |
| <u>* double (-bis) derivatization</u> |  |  |  |  |  |

### 2.2 Other chemicals and supplies

Derivatization of estrogens and estrogen metabolites were achieved with dimethylaminoaphthalene Dansyl Chloride ≥ 99.0% HPLC sourced from Sigma Aldrich (St. Louis, MO). Dansyl Chloride-D6 was sourced from C/D/N Isotopes, Inc. (Pointe-Claire, Quebec, Canada) and used to generate in-house internal standards for all 15 estrogens and estrogen metabolites. β-glucuronidase/sulfatase from Helix pomatia was obtained from Millipore Sigma (St. Louis, MO). Glacial acetic acid (Certified ACS), L-ascorbic acid (Certified ACS), acetone (HPLC grade), acetonitrile (Optima HPLC grade), and methanol (Optima HPLC grade) were purchased from Fisher Chemical (Pittsburgh, PA). Sodium bicarbonate is purchased from Sigma Life Science (S5761). Sodium carbonate enzyme grade (Anhydrous) was purchased from Fisher Scientific (Waltham, MA). Sodium acetate (for HPLC) was acquired from Honeywell / Fluka (Morris Plains, NJ). Supported liquid extraction 400uL cartridges, 1cc cartridges, and 2cc cartridges were acquired from Biotage® (Charlotte, NC). Mass spectrometry vials 0.3mL and 1mL were purchased from Thermo Scientific (Waltham, MA).

### 2.3 Patient participants

Free and total estrogens were determined in plasma and stool from healthy control participants. Participants included 10 males, 10 post-menopausal females, and 10 premenopausal females. All premenopausal females provided paired stool samples collected in both the follicular and luteal phases. Each premenopausal female also provided one plasma sample collected at the same time as one of the two stool samples. Thus, for each premenopausal participant, the stool samples were both follicular and luteal, while the plasma sample was either follicular or luteal (n = 5 for each plasma group). This study received approval from the institutional review boards at the University of California, Los Angeles.

### 2.4 Fresh frozen fecal sample collection

Healthy control participants were supplied with home stool collection kits and instructed to preserve their stool samples in a freezer immediately after collection until they could either be delivered to a study coordinator within 24–48 hours of collection. Participants were asked to provide stool consistent with their normal stool pattern and to avoid providing samples if they were constipated or had diarrhea. Subsequently, study coordinators transferred the samples to laboratory freezers for long-term storage at −80C.

### 2.5 Plasma sample collection

30 healthy control patients were instructed to come in for plasma collection. Plasma was collected through a plasmapheresis machine in heparinized tubes and spun down to remove cells. Patient plasmas were then stored in −80C until further sample processing.

### 2.6 Sample preparation optimization

Sample preparation procedures of plasma and stool samples were adapted from Xu (2005). Various experimental parameters were optimized through extraction efficiency.

#### 2.6.1 Basic reaction buffer optimization

Volume of sodium acetate buffer (0.15M) was optimized to 300 uL added to plasma samples. Volume of buffer was optimized to 1 mL in stool samples.

#### 2.6.2 L-ascorbic acid optimization

Study comparisons of 1 mg/mL, 5 mg/mL, and 10 mg/mL L-ascorbic acid solubilized in sodium acetate buffer (0.15M) yielded optimal extraction efficiency to 1 mg/mL.

#### 2.6.3 Sample modifier optimization

A comparison study of 0uL, 20uL, or 40uL of isopropanol (IPA) added to plasma samples yielded the best extraction efficiency for 20uL. Concentrations of 0%(v/v), 10%(v/v), 33%(v/v), and 50%(v/v) acetonitrile were compared to optimize extraction efficiency and found 33% ACN yielded optimal extraction efficiency.

#### 2.6.4 Enzyme optimization

15 uL of β-glucuronidase/Arylsulfatase enzyme was optimized for both plasma and stool samples after assessing 0, 10uL, 20uL, and 40uL of enzyme injection.

#### 2.6.4 Organic solvent optimization

A comparison study of extraction efficiencies organic solvents as eluents included dichloromethane (DCM), methyl tert-butyl ether (MTBE), and ethyl acetate (EtAc) were analyzed. DCM yielded the best extraction efficiency.

#### 2.6.5 Elution optimization

Efficiency of the number of organic solvent elutions was optimized after comparing 3x elutions (1.5 mL DCM each elution) vs. 5x. 3x yielded no significant difference between 5x, thus 3x elutions were optimal.

### 2.6 Plasma sample preparation

Plasma was prepared with and without enzymatic hydrolysis. For determination of free estrogen, 100 uL plasma was combined with stable heavy isotope internal standards and 300 uL of basic reaction buffer consisting (BRB) consisting of 0.15 M acetate buffer pH 4.5 containing 1.0 mg/mL L-ascorbic acid. For determination of total estrogen, 100 uL plasma was prepared as above, but 15 uL of β-glucuronidase/aryl-sulfatase (from Helix pomatia) was also added and samples were kept gently rocking at 37°C overnight. For both free and total samples, isopropanol was added (5% v/v) and lipids were extracted with 400 cc supported liquid extraction (SLE) cartridges (Biotage LLC) using 3 x 1.5 mL dichloromethane extractions and dried under argon.

### 2.8 Stool sample preparation

Stool for both free and total estrogen (100 mg each) was combined with internal standards and 1 mL BRB and subjected to bead mill homogenization (Biotage Lysera). Stool for total estrogen was combined with 15 uL deconjugation enzyme and reacted overnight as above. Both sample sets were combined with 500 ul acetonitrile; re-homogenized; and centrifuged. Supernatant was loaded into 2 cc SLE cartridges (Biotage) and extracted with 3 x 2 mL DCM followed by drying under argon.

### 2.9 Derivatization and derivatization optimization

Derivatization conditions were optimized for pH, optimal concentration of DS (mol ratio), temperature optimization, and duration (time) optimization. Optimized conditions were assessed using yield. A pH at 9.2 was found to be optimal. Heating sample at 60 degrees for 20 minutes resulted the best yield of dansylation, as overheating caused degradation of dansyl. There were no significant differences in yield between 10 minutes, 20 minutes, and 30 minutes. Optimized conditions for derivatization were as follows: 100 uL of 1mg/mL DSC in acetone with 100 uL of 0.1M sodium bicarbonate buffer (pH at 9.2). Samples were vortexed for 30 seconds and then incubated for 20 minutes at 60 degrees Celsius. Samples were resuspended to 200 uL with acetone and centrifuged. 100 uL of supernatant was transferred to 0.3mL MS vials. Reference supplemental figure for full workflow.

### 2.10 Estrogen DS-D6

E-DS-D6 functions as an analyte-specific error correction from plasma and stool matrix effects. In-house generated internal standards for all 15 estrogens and estrogen metabolites were synthesized from estrogen calibrants and dansyl chloride-d6 powder. Stored as a combined stock solution in −80C. 1 uL of analyte-specific stock solution IS was injected into samples immediately prior to ESI LC-MS/MS run.

### 2.11 LC-MS/MS

LC-MS/MS analysis was performed on an Agilent 1290/SCIEX QTrap 5500 system using principles of tuning, method validation, quality control, calibration, and quantitation that we previously described 4. MS/MS settings for derivatized estrogens and stable heavy isotope internal standards were determined using reference calibration and internal standards (Steraloids); values were comparable to those previously reported 2, 3. Liquid chromatography was performed using a Kinetex C18 1.7 um 2.1 mm x 150 mm column (Phenomenex), while the gradient transitioned from 90% water/0.1% formic acid to 95% acetonitrile/0.1% formic acid across 20 minutes. Concentration of all estrogens was determined as ng/mL plasma or ng/mg stool.

### 2.12 Shotgun metagenomic analysis

Fecal samples were collected in ethanol and then later aliquoted and sent to One Codex in a DNA stabilization buffer and sequenced on their commercial standard depth platform for shotgun sequencing. The average sequence depth per sample was 1,659,737 with a range between 113,671 to 6,568,260 paired reads. Sequencing was performed using the shotgun sequencing method at a depth of 2 million 2×150 bp read pairs. The resulting sequencing data were uploaded to One Codex’s analysis software and compared against a comprehensive database of over 127,000 microbial reference genomes. Several statistical post-processing steps by One Codex were implemented to eliminate false positives from contamination or sequencing artifacts.

The DNA extraction was carried out using the Qiagen DNeasy 96 PowerSoil Pro QIAcube HT extraction kit, and library preparation was done using the KAPA HyperPlus library preparation protocol. Sequencing was conducted on an Illumina NextSeq 2000 instrument. Raw sequencing data (FASTQ files) were analyzed using the One Codex database, which includes approximately 148,000 complete microbial genomes, covering bacteria, viruses, archaea, and eukaryotes. Human and mouse genomes are also included to filter out host reads. The metagenome annotation involved three steps: k-mer based classification using k-mers where k=31, artifact filtering to remove probable sequencing or reference genome artifacts, and species-level abundance estimation based on sequencing depth and coverage across reference genomes.

### 2.13 Statistics

Significance defined as p<0.05. For comparison with more than two groups: two way ANOVA with Sidak’s or Tukey’s multiple comparisons test and adjusted P values. For repeated measure comparison between two groups: paired Student’s t-test. Correlation was determined on the basis of the Pearson r for the linear regression and the P value for the significance of the correlation. Both r and P value were determined using correlation analysis within Prism.

## 3. Results

### 3.1 Development and validation of LC-MS/MS method for determination of free and conjugated primary estrogens and estrogen metabolites in human stool and plasma

Aromatase enzymes located in ovaries and non-ovarian tissue synthesize estrone and estradiol from the androgens androstenedione and testosterone. Estrone and estradiol are subsequently metabolized by CYP450 mediated phase I hydroxylation and COMT-mediated phase II methoxylation (**Figure 1A**). There exist validated LC-MS/MS methods for determining the levels of these primary estrogen and estrogen metabolites in human plasma, serum, and urine. However, there exist no reports of a fully validated method for the determination of estrogens in human stool; nor any determination of human stool estrogens together with human serum or plasma estrogens. We thus sought to overcome this limitation in prior research by developing and validating an LC-MS/MS method for determining in human stool and plasma the levels of 15 estrogen and estrogen metabolites (**Table 1**).

We sought to determine estrogen levels in the stool and plasma of human males and pre- and postmenopausal females. We thus required a highly sensitive LC-MS/MS method capable of detecting estrogens across the reference ranges for males and postmenopausal women. It is often possible to measure small molecules by LC-MS/MS without further chemical modification. However, unmodified estrogen ionizes poorly in LC-MS/MS, separates with poor resolution in standard LC, and yields limits of detection well outside these reference ranges [34].

The derivatizing agents dansyl chloride (DSC) and pyridine-3-sulfonyl chloride are commonly employed to increase sensitivity of estrogen detection in LC-MS/MS [31]. We evaluated pyridine-3-sulfonyl chloride but settled upon dansyl chloride under standard reaction conditions [38] because of the avidity and simplicity of its derivatization reaction (**Figure 2A**). We determined optimum MS/MS settings for all 15 analytes and two class specific stable heavy isotope internal standards by tuning upon reference calibrants (**Figure 2B** and **Supplemental Table 1**). Of note, MS/MS alone is incapable of fully resolving isobaric estrogens—those that share a molecular weight. For example, 2-MeOE1, 3-MeOE1, and 4-MeOE1 all have the same monoisotopic mass (**M_min_**), as do their dansyl (DS) derivatives; and the DS species likewise share parent and fragment ions. Chromatography must thus be used to separate isobaric analytes. We tested a number of LC columns and gradient conditions including Kinetex C8, Evo C18 1.7 um, and biphenyl columns, but we achieved the best resolution using a Kinetex C18 1.7 um 2.1 mm x 150 mm column and an acetonitrile/0.1% formic acid gradient (**Figure 2C**). These LC conditions successfully resolved isobaric species including 2/3/4-MeOE1 (**Figure 2C**). Final calibration curves had excellent linearity (**Figure 2D**) across 4 orders of magnitude, and limits of detection and quantitation for all analytes were approximately 1 pg/mL (**Table 2**).

**Figure 2.**
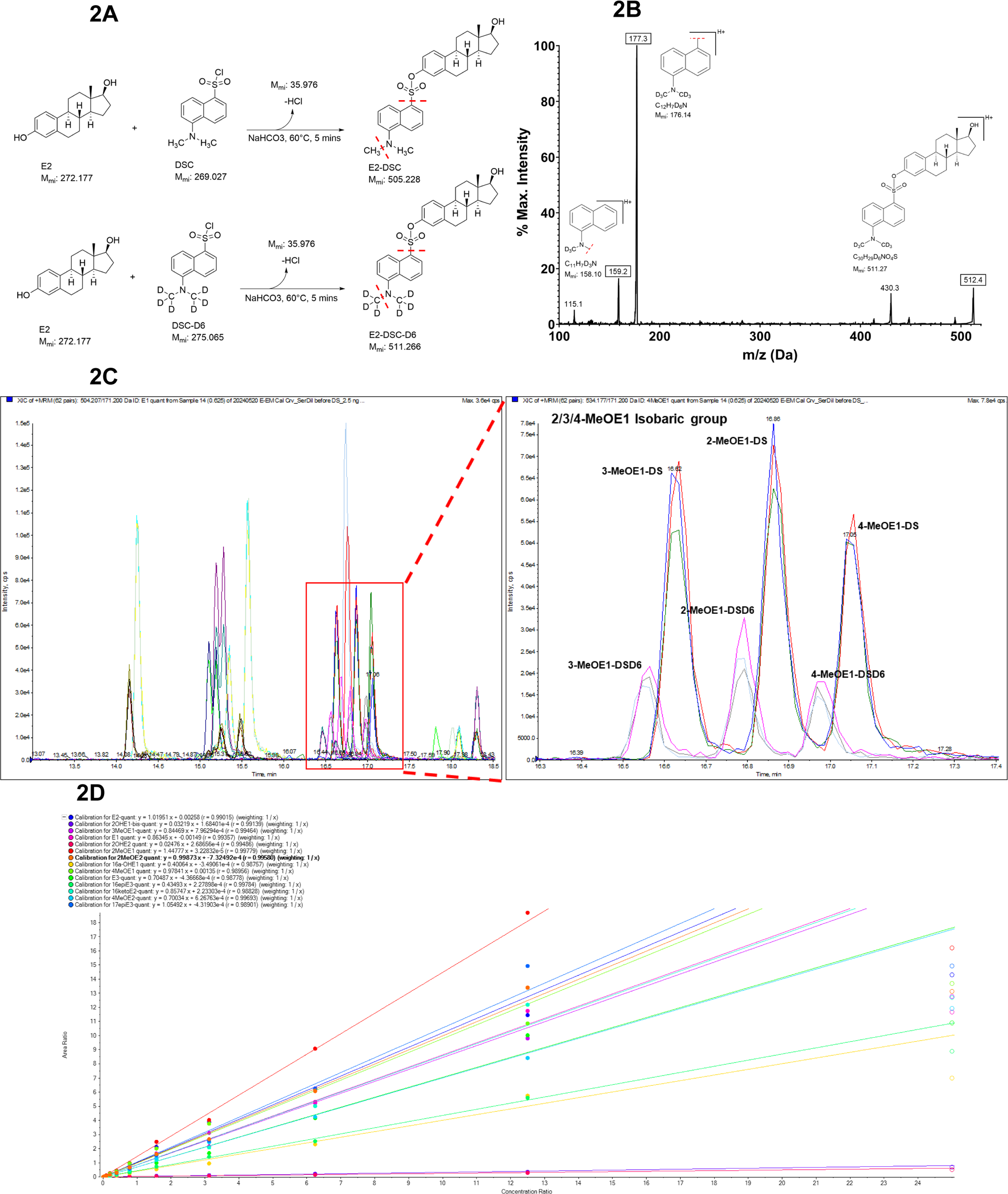
Derivatization, mass spectrometry, and liquid chromatography of estrogens. (**A**) Reaction mechanism for derivatization of estrogen with DSC and DSC-D6. Red dashed lines: MS/MS fragmentation breaks for MS2 ions detected in LC-MS/MS method. (**B**) Estradiol together with its MS/MS fragmentation pattern. Ions employed within quantitative and qualitative transitions are given as structural diagrams and [M+H]. (**C**) Liquid chromatogram of all E-DS and E-DS-D6 peaks. (*Inset*) Representative isobaric groups at 534.2 (2/3/4-MeOE1-DS) and 540.2 (2/3/4-MeOE1-DS-D6) are successfully resolved by LC. Abbreviation: **M_mi_**, monoisotopic mass.

**Table 2.** Linearity and linear range of detection for estrogen LC-MS/MS analytes. **LLOD**, lower limit of detection; **LLOQ**, lower limit of quantitation; **ULOQ**, upper limit of quantitation.

|  | **LLOD (pg/ml)** | **LLOQ (pg/ml)** | **ULOQ (pg/mL)** | **r** |
| --- | --- | --- | --- | --- |
| Estradiol (E2) | 0.38 | 1.5 | 12500 | 0.99 |
| 2-OHE1 | 6.1 | 6.1 | 25000 | 0.99 |
| 3-MeOE1 | 0.38 | 0.76 | 12500 | 0.99 |
| Estrone (E1) | 0.76 | 1.5 | 12500 | 0.99 |
| 2-OHE2 | 6 | 12.2 | 25000 | 0.99 |
| 2-MeOE1 | 0.38 | 0.76 | 12500 | 0.99 |
| 2-MeOE2 | 0.38 | 1.5 | 12500 | 0.99 |
| 4-OHE1 | 6 | 6 | 25000 | 0.99 |
| 4-MeOE1 | 0.38 | 1.5 | 12500 | 0.99 |
| 16α-OHE1 | 0.38 | 1.5 | 12500 | 0.99 |
| 4-MeOE2 | 0.76 | 1.5 | 12500 | 0.99 |
| 17-epiE3 | 0.38 | 1.5 | 12500 | 0.99 |
| Estriol (E3) | 0.38 | 1.5 | 12500 | 0.99 |
| 16-epiE3 | 0.38 | 0.76 | 25000 | 0.99 |
| 16-ketoE2 | 0.38 | 1.5 | 12500 | 0.99 |

Estrogens are present in the circulation and stool as both free species and sulfate or glucuronide conjugates (**Figure 1B**). It is possible, however, to deconjugate estrogen during sample preparation by addition of beta-glucuronidase and arylsulfatase enzyme [38]. We thus developed sample preparation methods for determination of both free and total (free + conjugated) estrogen in plasma and stool. We modified a sample preparation method previously validated for serum [37] but optimized all parameters for plasma and stool. Thus, we compared the plasma estrogen extraction efficiency of liquid-liquid extraction (LLE) versus supported liquid extraction (SLE), and across the extraction solvents dichloromethane (DCM), methyl-tert-butyl-ether (MTBE) and ethyl acetate (EtAc) (**Figure 3A-B**). SLE with DCM had both greater throughput and superior extraction efficiency. We also optimized stool extraction conditions for SLE (**Figure 3C**) and determined that bead mill homogenization was superior to hand held homogenization (**Figure 3D**).

**Figure 3.**
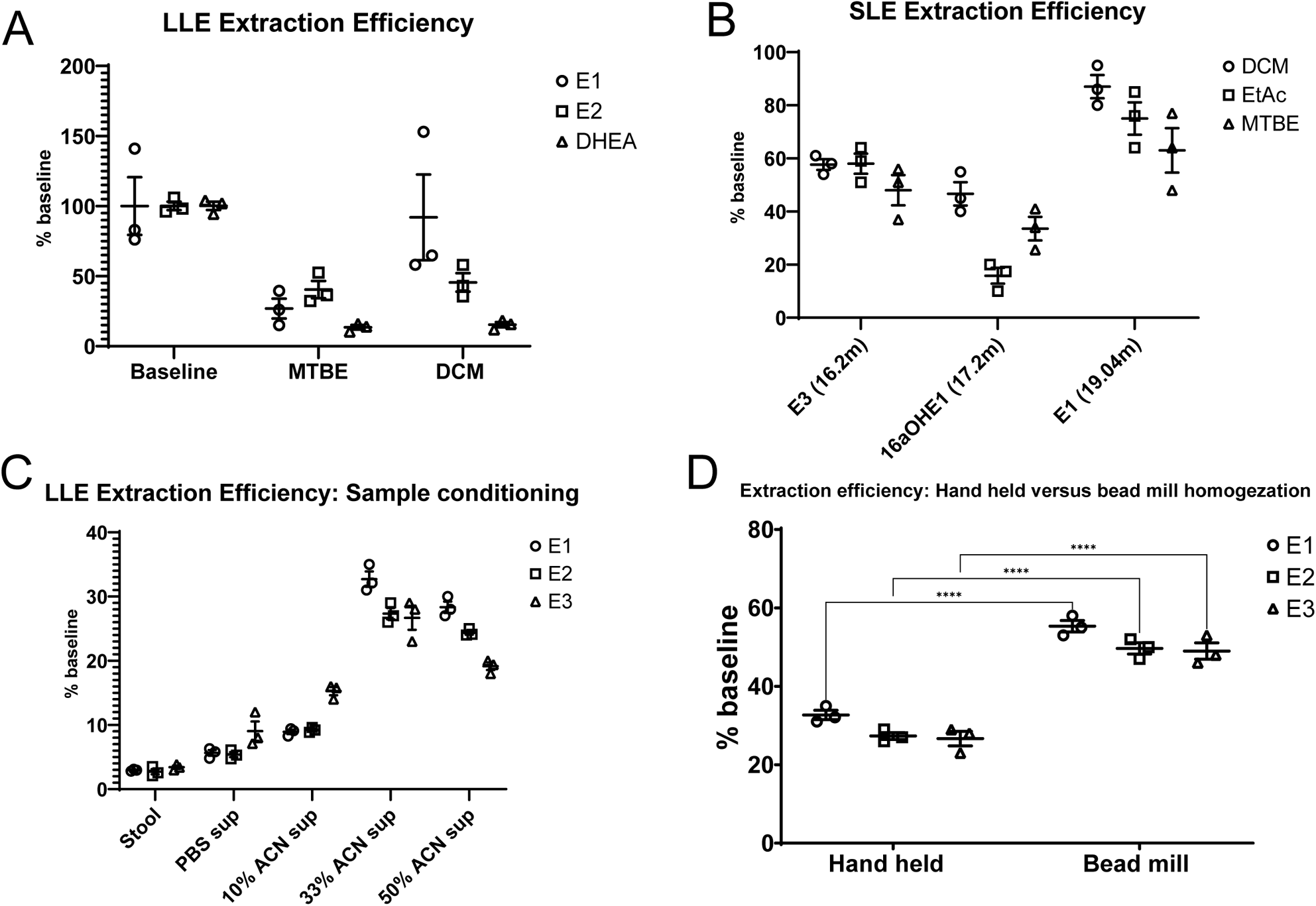
Representative optimizations of plasma and stool sample preparation methods. (**A**) LLE extraction efficiency for representative estrogens given compared to baseline, for both MTBE and DCM. (**B**) SLE extraction efficiency for representative estrogens given compared to baseline, for MTBE, DCM and EtAc. (**C**) LLE extraction efficiency determined for different stool sample conditioning methods, following hand held homogenization of stool samples. (**D**) Hand held homogenization compared to bead mill homogenization with respect to extraction efficiency of representative estrogens. (Statistics: ****, P < 0.0001. Two-way ANOVA with Šídák’s multiple comparisons test and adjusted P values.)

Plasma and stool exhibits considerable differential matrix effect—the effect that co-eluting material from the original sample has upon ionization efficiency (**Figure 4A-B**). The consequence of differential matrix effect is to introduce inaccuracy into concentration determination and across samples—as an analyte whose ionization is not suppressed in the calibration curve gets variably suppressed by differential matrix both intra- and inter-sample. Assuming the samples are of uniform rather than variable matrix, it is possible to overcome this problem by building calibration curves in a representative matrix. It is also possible to correct for matrix effect by employing analyte-specific co-eluting heavy stable isotope labeled internal standards [40]. However, while charcoal stripped plasma can serve as a representative plasma matrix, given the manifest inter-stool variability, there exists no such representative matrix for stool. At the same time, It is difficult and costly to acquire stable isotope labeled estrogens for all 15 estrogen analytes of the present method. Because of these constraints, we therefore synthesized our own analyte-specific heavy stable isotope labeled internal standards from both estrogen calibrants and heavy stable isotope labeled dansyl chloride (DS-D6) (**Figure 2A**). These DS-D6 internal standards, added directly the final MS sample, co-elute with the estrogen analytes (**Figure 2C**) and can be used to correct for matrix effect. In combination with a class specific internal standard added at the beginning of sample preparation, both technical loss and matrix effect can be successfully corrected for (**Figure 2D**).

**Figure 4.**
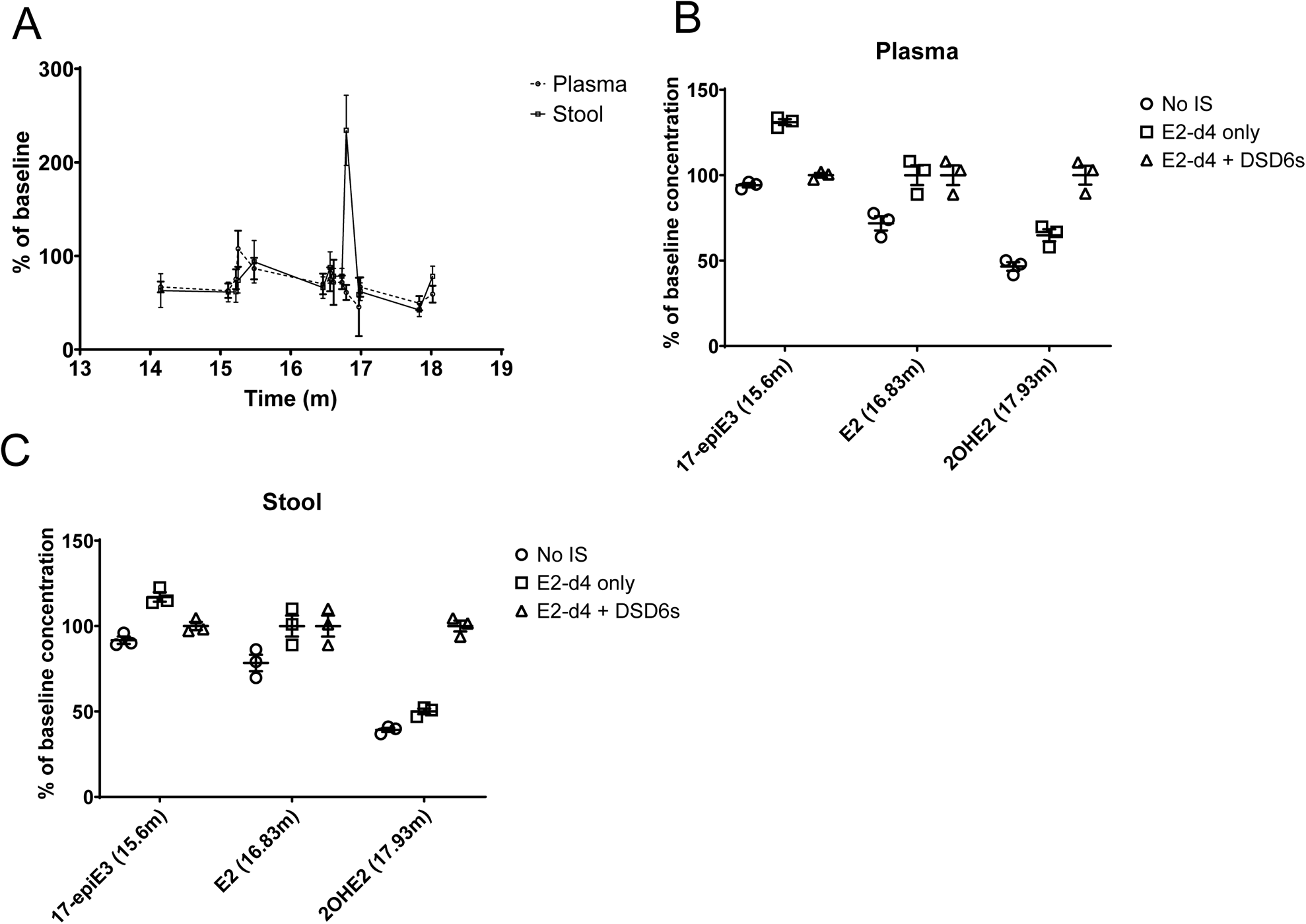
Use of analyte specific DSD6 internal standards overcomes matrix effect and rescues accuracy of estrogen LC-MS/MS. (**A**) Class specific E2-D4 internal standard was to charcoal stripped plasma or uniform stool matrix; samples were extracted and derivatized; and analyte specific DSD6 was added in final MS sample. Effect of matrix on DSD6 compared to no-matrix baseline was determined. (**B-C**) 17epiE3, E2, and 2OHE2 were spiked into the plasmas (B) and stools (C) of Fig4A. Concentration was determined without use of IS; with class-specific correction only; and with class-specific correction combined with analyte-specific correction for matrix effect.

Our final sample preparation workflows for determination of free and total estrogen in human plasma and stool are represented herein (**Figure 3**). We determined accuracy and precision for plasma and stool determination of free estrogen by spiking low, medium, and high concentrations of all 15 estrogens into charcoal stripped plasma or a uniform stool matrix (**Supplemental Table 2**). RSD across 5 replicates averaged from between 13% and 19% for plasma at each of the spike concentrations. RSD for stool ranged from 14% to 21% for all analytes at all concentrations. Accuracy was acceptable with skew of <20% across most all conditions.

**Figure 5.**
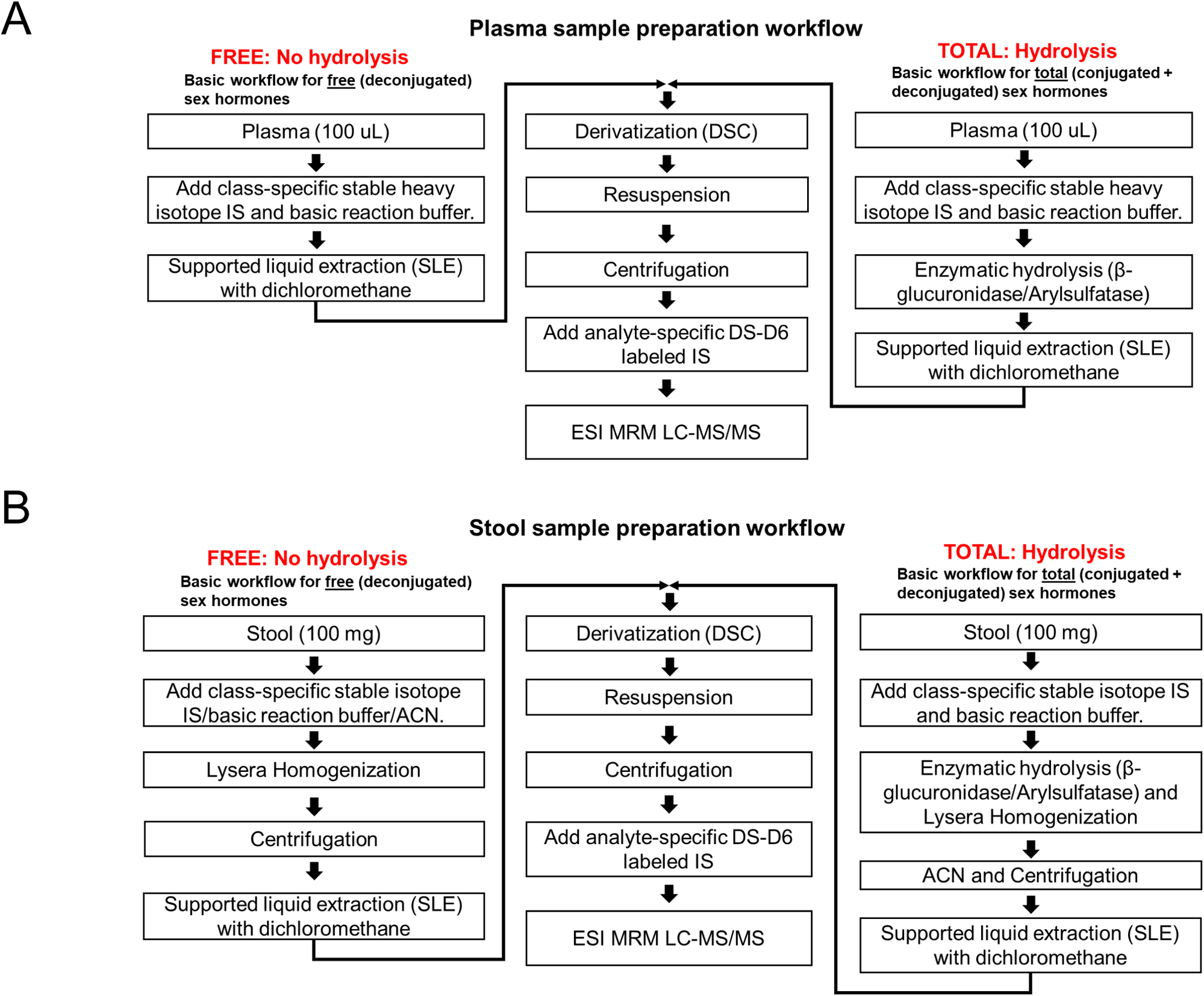
Overall sample preparation workflows for plasma and stool. (A) Sample preparation workflow for plasma. (B) Sample preparation workflow for stool.

### 3.2 Determination of free and total estrogens in human plasma and stool resolve differences in sex, menstrual cycle, and menopausal status

We gathered plasma and stool from a small cohort of healthy control human subjects. Study participants included 10 males, 10 post-menopausal females, and 10 premenopausal females. All premenopausal females provided paired stool samples collected in both the follicular and luteal phases. Each premenopausal female also provided one plasma sample collected at the same time as one of the two stool samples. Thus, for each premenopausal participant, the stool samples were both follicular and luteal, while the plasma sample was either follicular or luteal (**Figure 6A**).

**Figure 6.**
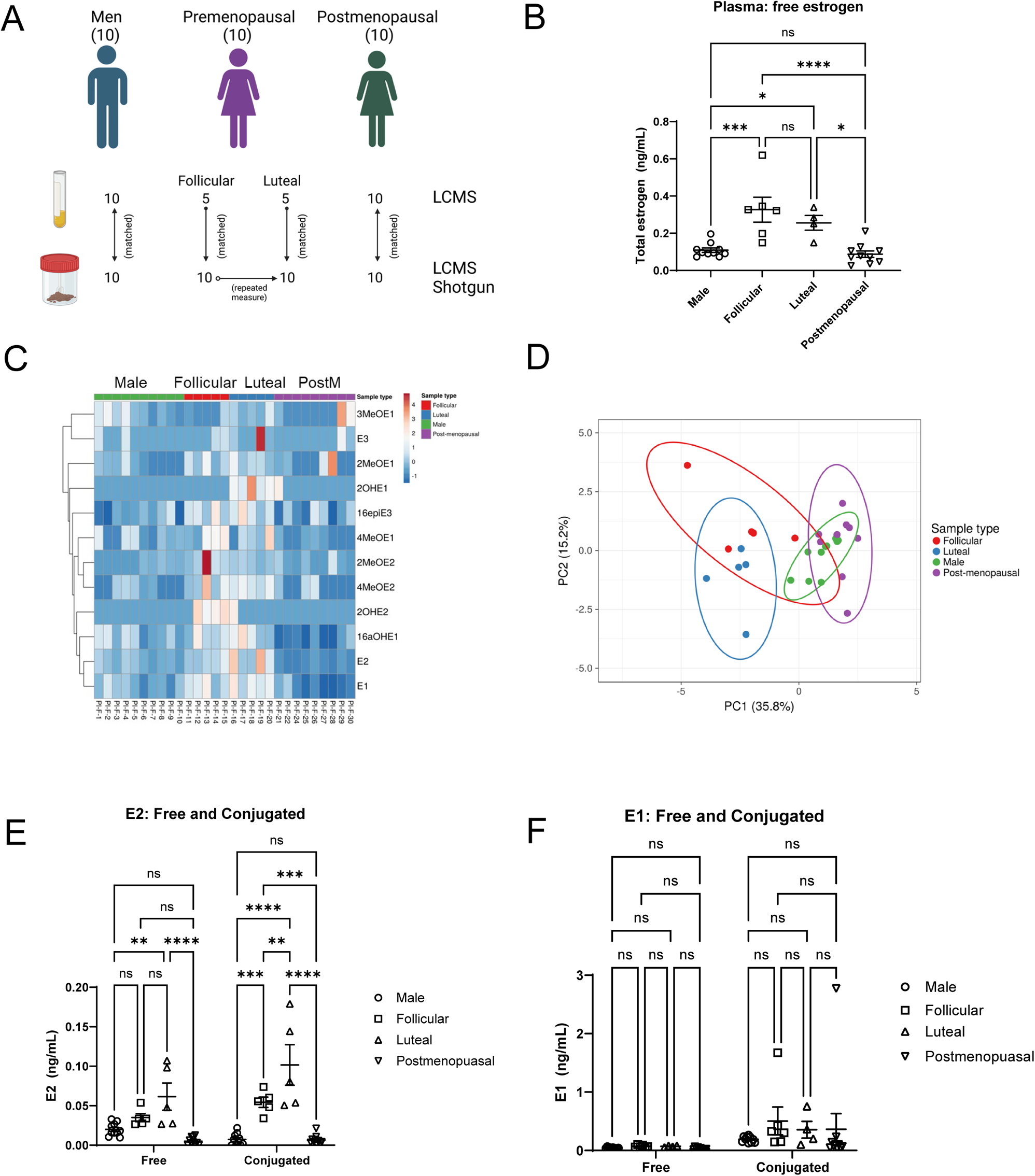
Determination of free and total estrogen in plasma of healthy control human study participants resolves differences between sex, menstrual cycle, and menopausal status. (**A**) Study design. (**B**) The sum of all free estrogens within a sample was compared across groups. (**C**) Heatmap of free estrogen within each sample, with clustering by analyte. (**D**) Unit variance scaling is applied to rows; SVD with imputation is used to calculate principal components. X and Y axis show principal component 1 and principal component 2 that explain 35.8% and 15.2% of the total variance, respectively. Prediction ellipses are such that with probability 0.95, a new observation from the same group will fall inside the ellipse. N = 29 data points. (**E-F**) The concentrations of free and conjugated E2 (E) and E1 (F) were determined for all groups. *Statistics*: *, P < 0.05; **, P < 0.01; ***, P < 0.001. Two-way ANOVA with Tukey’s multiple comparison test and adjusted P values.

We first determined the levels of free and total estrogens in the plasma from this study. We observed that both follicular and luteal plasma had significantly higher levels of all free estrogens taken together than did men and postmenopausal women (**Figure 6B**). A heatmap comparing the individual free estrogens reproduced this pattern (**Figure 6C**), while dimensional reduction of the collective free estrogen data clearly resolved differences between not only men and pre- and postmenopausal women but between follicular and luteal phase as well (**Figure 6D**). Comparing free and conjugated levels of plasma estrogen revealed that free E2 was significantly elevated in luteal plasma while conjugated E2 was significantly higher in both follicular and luteal plasma, while postmenopausal women produced almost no E2 (**6E**). By contrast, there were no significant differences across all groups with respect to both free and total E1 (**6F**). Furthermore, while E2 was present equally as both free and conjugated, E1 was largely present as conjugated in all groups.

We also determined the levels of free and total estrogens in the stool from all healthy control participants. The sum all total (ie free + conjugated) estrogens were significantly elevated in luteal stool (**Figure 7A**). Consistent with this result, heatmap of total estrogen for each analyte reveals elevation in luteal stool (**Figure 7B**), while dimensional reduction clearly resolves this group (**Figure 7C**). When comparing the sum of all total (free + conjugated) estrogens in repeated measure follicular versus luteal, both the sum of total estrogen (**Figure 7D**) and total estradiol (**Figure 7E**) significantly increased across the menstrual cycle.

**Figure 7.**
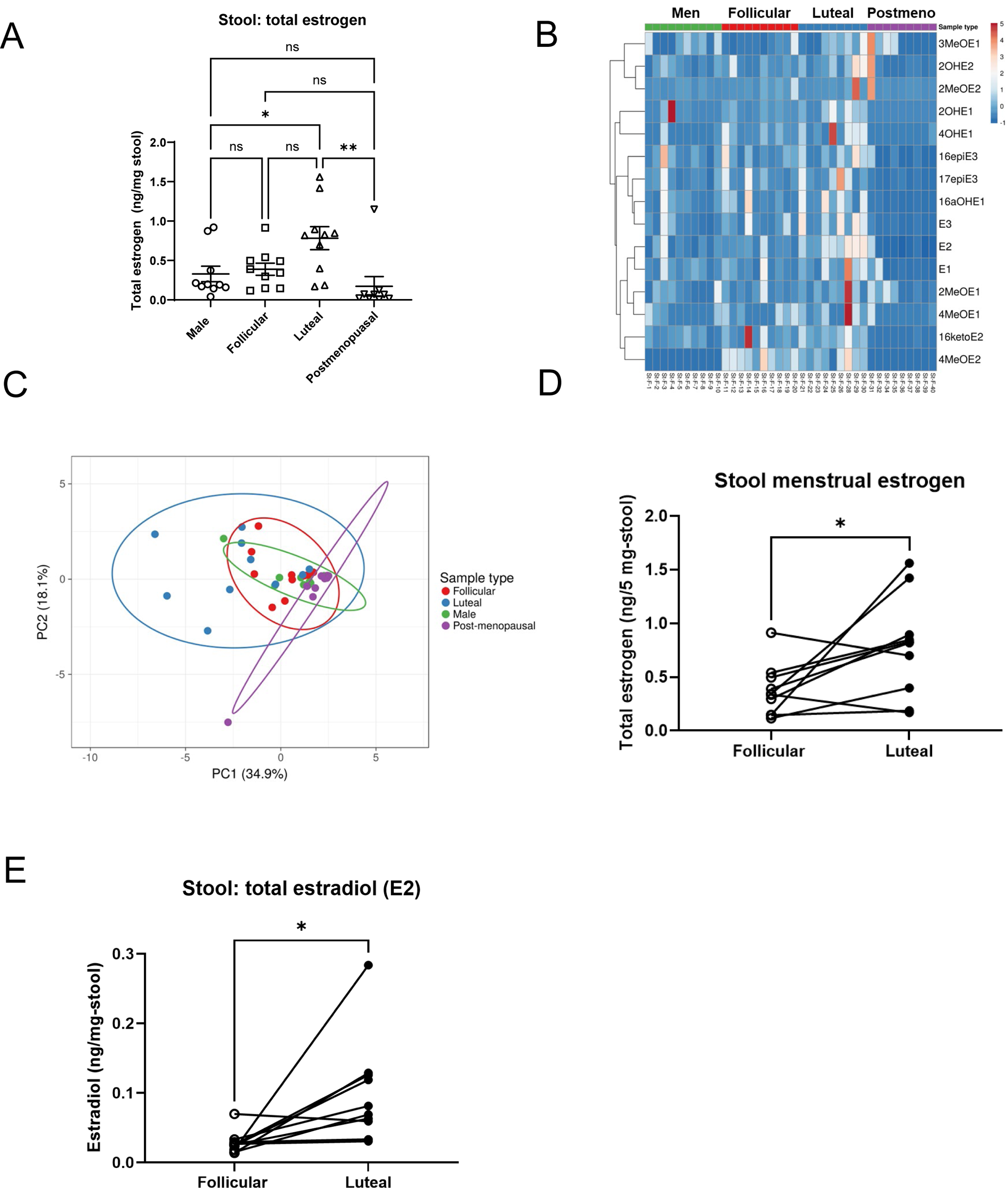
Determination of free and total estrogen in stool of healthy control human study participants resolves differences between sex, menstrual cycle, and menopausal status. (**A**) Free and total (free + conjugated) estrogen levels were determined for each analyte in the stool of all study participants. The sum of all total (free + conjugated) estrogen analytes was compared across all groups. (**B**) Heatmap of total (free + conjugated) estrogen analytes within each sample, with clustering by analyte. (**C**) Unit variance scaling is applied to rows; SVD with imputation is used to calculate principal components. X and Y axis show principal component 1 and principal component 2 that explain 35% and 18% of the total variance, respectively. Prediction ellipses are such that with probability 0.95, a new observation from the same group will fall inside the ellipse. N = 38 data points. (D) The sum of all total estrogen analytes was compared in matched follicular and luteal stool samples. (E) The sum of total estradiol was compared in matched follicular and luteal stool samples. *Statistics*: *, P < 0.05; **, P < 0.01; ***, P < 0.001. For comparisons between more than two groups: two-way ANOVA with Tukey’s multiple comparison test and adjusted P values. For comparison between repeated measure: Paired Student’s t-test.

### 3.3 Correlations between plasma and stool estrogen suggest that intestinal excretion of estrogen is dependent on systemic concentration

We determined the correlations between plasma and stool estrogen, comparing the sum of all total (free + conjugated) estrogen analytes in both plasma and stool (**Figure 8A**) as well as representative analytes including total estrone in plasma versus stool (**Figure 8B**). We observed that plasma total estrogen strongly and highly significantly correlated with stool total estrogen, as did individual analytes including estrone. These results are consistent with the rate of estrogen excretion into stool being driven by systemic estrogen concentration.

**Figure 8.**
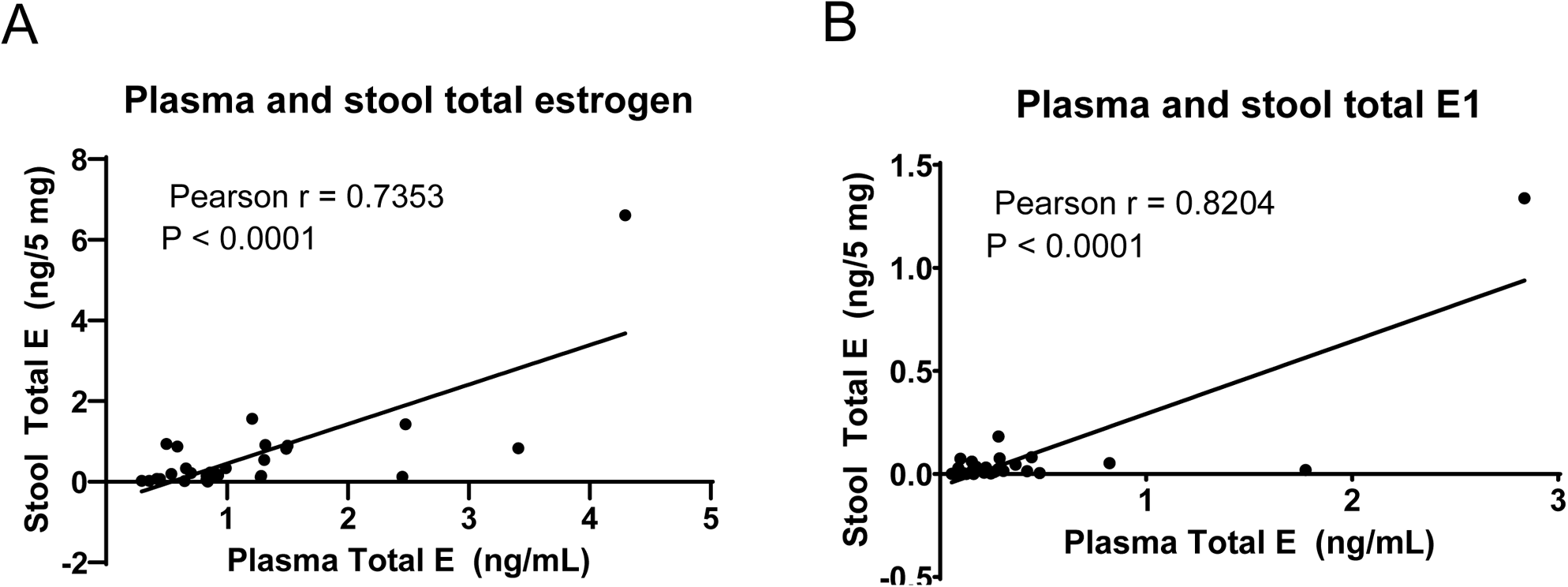
Total estrogen level in plasma is highly correlated with total estrogen level in stool. (**A**) Correlation between the sum of all total (free + conjugated) estrogen analytes in both plasma and stool was determined. (**B**) Correlation between total plasma estrone with total stool estrone was determined.

### 3.4 Shotgun metagenomic sequencing reveals sex differences in the functional capacity of microbiota to make estrogen reactivating enzymes

We subjected all stool samples to shotgun metagenomic sequencing, from which we extracted read depth-normalized gene copy number for the KEGG orthologs for β-glucuronidase (K01195) and arylsulfatase (K01130). We then compared the functional capacity to make both β-glucuronidase (**Figure 9A**) and β-glucuronidase + arylsulfatase (**Figure 9B**) between all four groups. We observed that these functional capacities to make sex hormone reactivating enzymes were significantly elevated in follicular phase stool.

**Figure 9.**
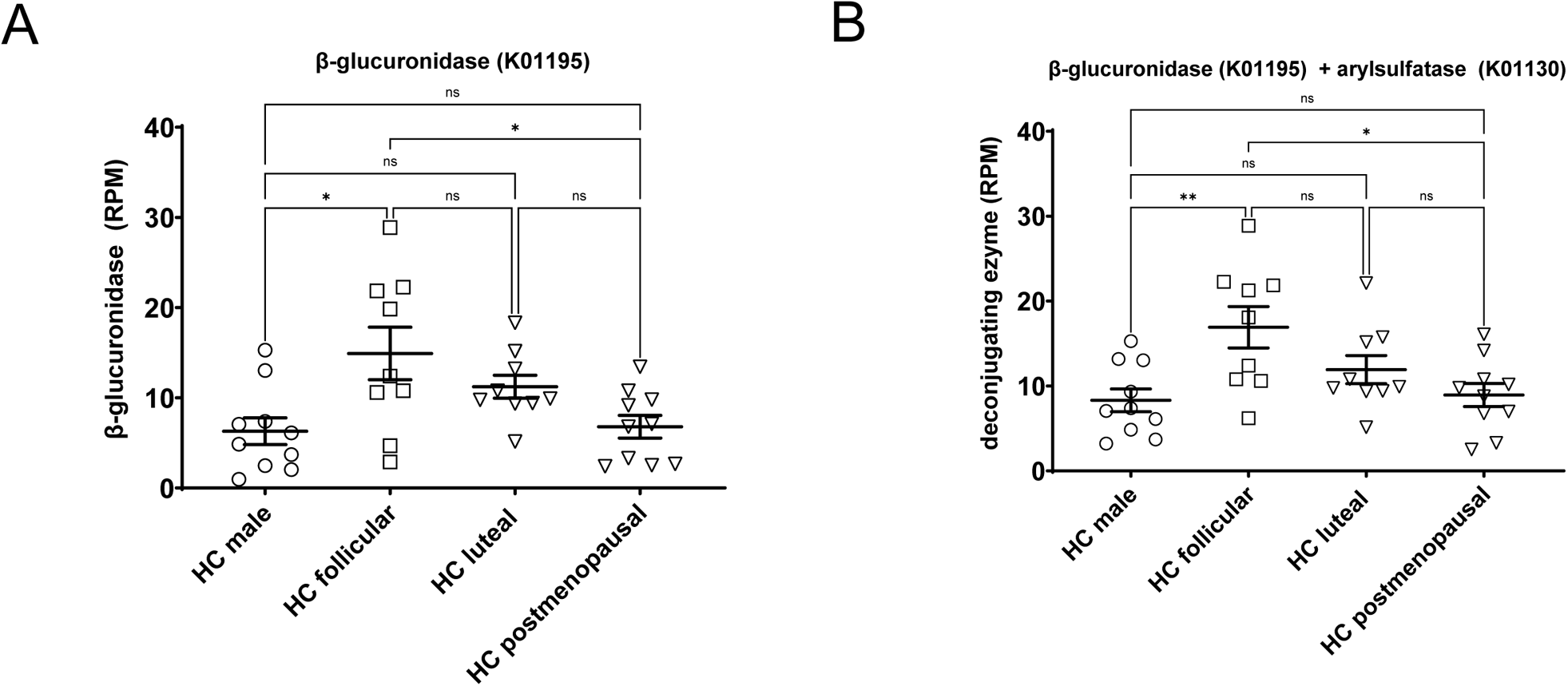
There exists a sex difference in the functional capacity of gut microbiota to produce the estrogen reactivation enzymes beta-glucuronidase and arylsulfatase. Stool samples were subjected to shotgun metagenomic sequencing and normalized gene copy number for both beta-glucuronidase (K01195) and arylsulfatase (K01130) were determined. Levels of functional capacity to produce beta-glucuronidase alone (**A**) and beta-glucuronidase + arylsulfatase (**B**) were compared across all groups. Statistics: Two-way ANOVA with Tukey’s multiple comparisons test and adjusted P values.

### 3.5 Shotgun metagenomic sequencing and LC-MS/MS analysis indicate that both beta-glucuronidase and arylsulfatase together, and not beta-glucuronidase alone, are responsible for reactivation of gut estrogens

We combined the LC-MS/MS and gene sequencing datasets in order to ask the question whether the gut microbial reactivation of stool estrogen was a function of one or both of microbial β-glucuronidase or arylsulfatase. We thus compared both beta-glucuronidase RPM alone and beta-glucurondiase + arylsulfatase RPM, with the amount of reactivated estrogen in each of the four groups. Interestingly, we observed that beta-glucuronidase alone failed to correlate with the amount of reactivated estrogen for each of the groups including males (**Figure 10A**) and follicular stool (**Figure 10B**). By contrast, beta-gluronidase and arylsulfatase RMP together positively correlated with reactivated estrogen in males (**Figure 10C**) and follicular stool (**Figure 10D**) while trending significant in both luteal and postmenopausal stool (data not shown).

**Figure 10.**
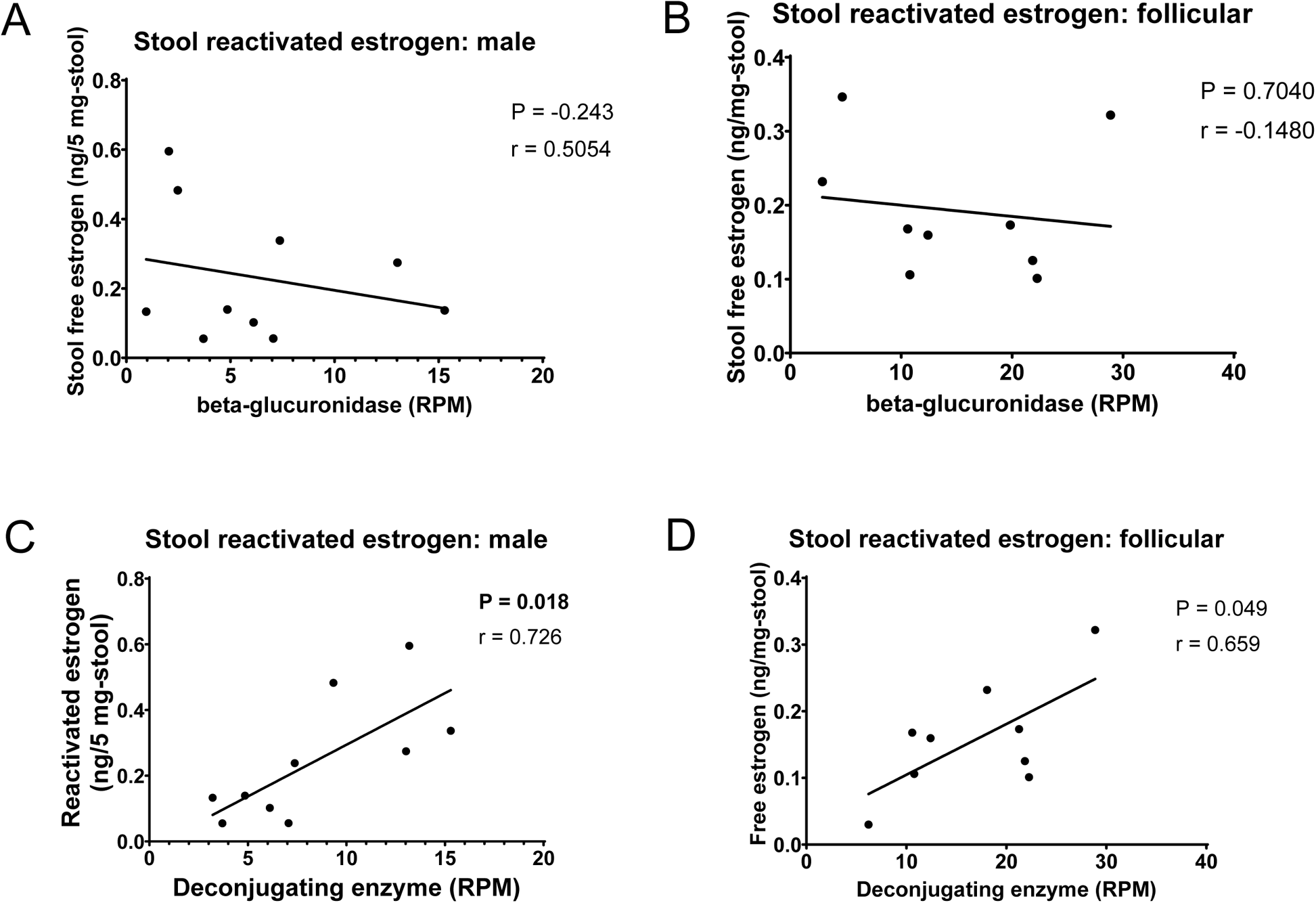
Beta-glucuronidase + arylsulfatase gene copy number, but not beta-glucuronidase gene copy number alone, correlated with reactivated estrogen in male and female stool. Gut microbial capacity to make beta-glucuronidase alone or beta-glucuronidase + arylsulfatase was correlated with the level of reactivated estrogen in all groups including male (**A**) and follicular stool (**B**). Correlation was determined as significance with respect to Pearson r in Prism software.

### 3.6 Shotgun metagenomic sequencing and LC-MS/MS analysis indicate that gut reactivation of estrogen might modulate systemic estrogen concentration

We also compared gene copy number of beta-glucuronidase + arylsulfatase with both the sum of all total (free + conjugated) plasma estrogen levels as well as individual total plasma estrogen levels. The sum of all total estrogens was correlated with deconjugation enzyme gene copy number in males (**Figure 11A**), while individual plasma estrogen analytes were correlated with deconjugation enzyme RPM in males (**Figure 11B**) and premenopausal women (**Figure 11C**).

**Figure 11.**
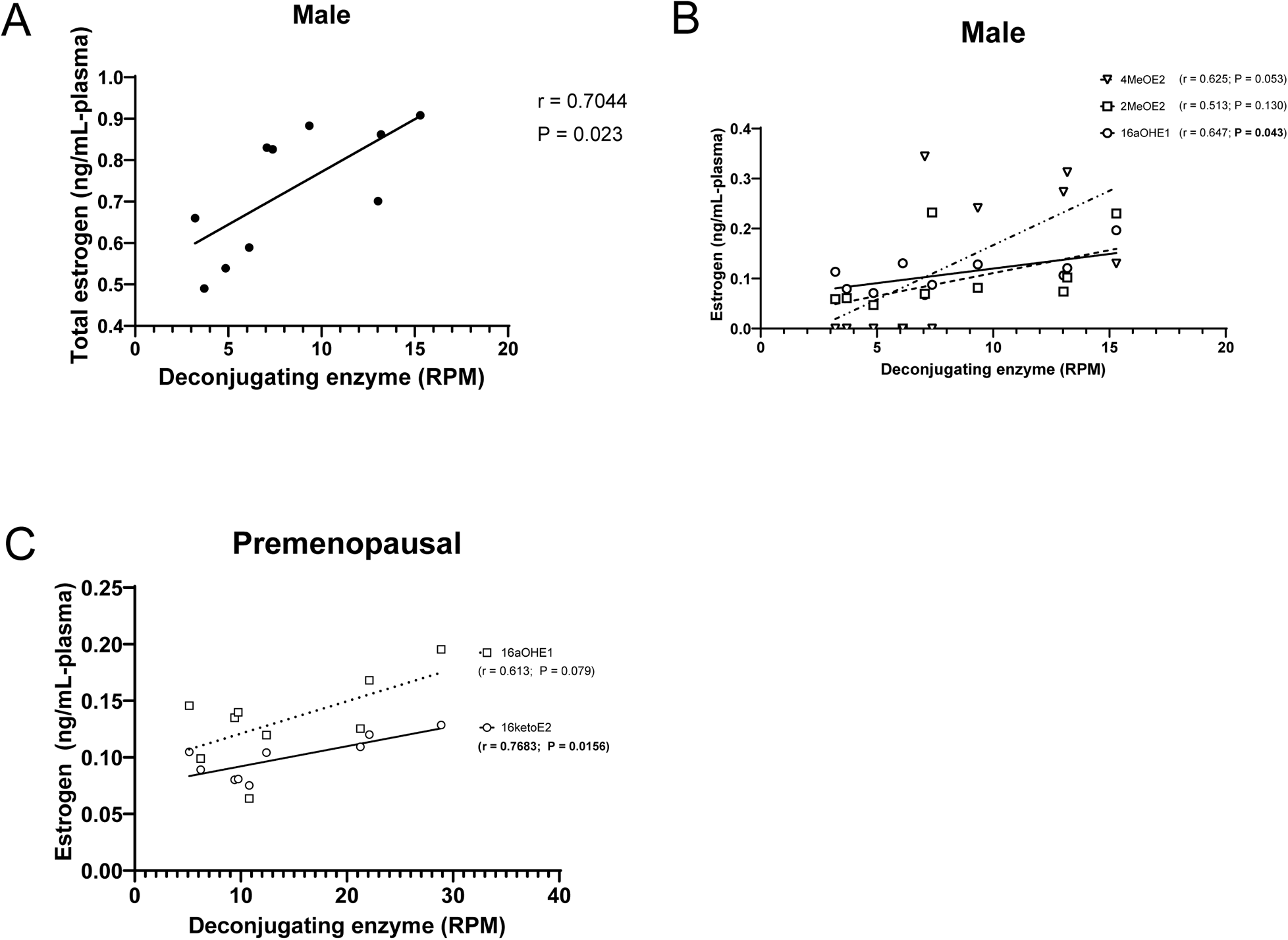
Gut microbial capacity to make beta-glucuronidase + arylsulfatase correlates with estrogen levels in male and female plasma. Beta-glucuronidase + arylsulfatase gene copy number was correlated with the sum of total (free + conjugated) estrogens in males (**A**) as well as individual total estrogens in males (**B**) and premenopausal females (**C**). Correlation was determined as significance with respect to Pearson r in Prism software.

## 4 Discussion

Human stool is a promising source of estrogen measurement, as it contains both free and conjugated forms of the hormone, as well as its metabolites, derived from both the host and microbiota. Stool samples are easy to collect, store, and transport, and they may provide a more stable and comprehensive indicator of estrogen exposure and metabolism than blood or urine. At minimum, stool estrogen measurement should be an important complement to systemic measures. Moreover, measurement of stool estrogen is necessary to address specific gaps in knowledge involving the relationship between systemic estrogen, stool estrogen, and estrogen reactivation in the gut.

It is surprising that there exist to our knowledge no validated LC-MS/MS methods for the determination of estrogens in human stool, and no published datasets combining stool estrogen with plasma or serum estrogen values. In order to fill this limitation in prior knowledge, we developed and validated an LC-MS/MS method for the determination of free and conjugated estrogens in stool and plasma. We employed derivatization techniques that had been established for serum estrogen, but we nonetheless optimized all aspects of sample preparation and LC-MS/MS. Of note, we employed SLE rather than LLE to extract estrogens from plasma and stool matrices. We further used high throughput bead homogenization to increase extraction efficiency from stool. And last, we synthesized stable isotope labeled analyte specific internal standards from DS-D6 that we used in combination with class specific internal standards to rescue the accuracy estrogen measurement from the matrix effects of both plasma and stool.

We had identified several knowledge gaps resulting from the limitation of prior knowledge that our present results at least partially address.

First, the mechanism by which the excretion rate of conjugated estrogens is determined is not clear. In response, we observed a strong positive correlation between systemic estrogen levels and levels in the stool (Figure 8). This result is consistent with excretion rate being driven directly by systemic concentration rather than by an additional circuit.

Second, it has not been clear whether gut microbial β-glucuronidase alone is largely responsible for reactivation of gut estrogen or whether arylsulfatase enzymes also play a role. We observed that only beta-glucuronidase + arylsulfatase gene copy number and not beta-glucuronidase alone correlated with reactived stool estrogen (Figure 10). This result strongly suggests that reactivation is a function of both enzymes, and that estrogen-sulfates make up a significant portion of conjugated estrogen excreted into the gut.

Third, there is little direct evidence for the hypothesis that gut microbial reactivated estrogen can be reabsorbed intestinally and thereby modulate systemic estrogen levels. We observed correlations between gut microbial deconjugation enzyme gene copy number and systemic estrogen levels in both men and premenopausal women (Figure 11). These results support that hypothesis that the gut microbiota can modulate systemic estrogen levels.

## Abbreviations

16-epiE3: 16-epiestriol
16-ketoE2: 16-ketoestradiol
16α-OHE1: 16α-hydroxyestrone
17-epiE3: 17-epiestriol
17β-HSD: 17β-Hydroxysteroid dehydrogenase
2-MeOE1: 2-methoxyestrone
2-MeOE2: 2-methoxyestradiol
2-OHE1: 2-hydroxyestrone
2-OHE2: 2-hydroxyestradiol
3-MeOE1: 3-methoxyestrone
4-MeOE1: 4-methoxyestrone
4-MeOE2: 4-methoxyestradiol
4-OHE1: 4-hydroxyestrone
COMT: catechol-o-methyltransferase
CYP19A1: cytochrome P450 family 19 subfamily A member 1
CYP450: cytochrome P_450_
DS: dansyl
DSC: dansyl chloride
E1: estrone
E1-d4: estrone-2,4,16, 16-d4
E2: estradiol
E2-d4: estradiol-2,4,16,16-d4
E3: estriol
ELISA: enzyme linked immunoassay;
IBS: irritable bowel syndrome
LC-MS/MS: liquid chromatography-mass spectrometry/mass spectrometry
LLOD: lower limit of detection
LLOQ: lower limit of quantitation
M_min_: monoisotopic mass
SULT: sulfotransferase
UGT: UDP-glucuronosyltransferases
ULOQ: upper limit of quantitation.

---

**Supplemental Table 1.** MS/MS settings together with retention times (RT) for all estrogen-DS and estrogen-DSD6 analytes.

| <u>Analyte</u> | <u>Mmi</u> | <u>Q1 mass</u><br><u>(Da)</u> | <u>DP (v)</u> | <u>EP (v)</u> | <u>Q3 mass</u><br><u>(Da)</u> | <u>CE (v)</u> | <u>CXP (v)</u> | <u>RT</u><br><u>(min)</u> |
| --- | --- | --- | --- | --- | --- | --- | --- | --- |
| <u>E1-DS</u> | <u>503.19</u> | <u>504.2</u> | <u>61</u> | <u>10</u> | <u>171</u> | <u>43</u> | <u>10</u> | <u>17.09</u> |
|  |  |  |  |  | <u>156</u> | <u>51</u> | <u>18</u> |  |
| <u>E2-DS</u> | <u>505.21</u> | <u>506.2</u> | <u>80</u> | <u>10</u> | <u>171</u> | <u>45</u> | <u>10</u> | <u>16.83</u> |
|  |  |  |  |  | <u>156</u> | <u>55</u> | <u>20</u> |  |
| <u>E3-DS</u> | <u>521.2</u> | <u>522.2</u> | <u>86</u> | <u>10</u> | <u>171</u> | <u>45</u> | <u>8</u> | <u>14.28</u> |
|  |  |  |  |  | <u>156</u> | <u>57</u> | <u>8</u> |  |
| <u>2-OHE1-bisDS</u> | <u>519.19</u> | <u>753.2</u> | <u>61</u> | <u>10</u> | <u>170</u> | <u>41</u> | <u>8</u> | <u>18.12</u> |
|  |  |  |  |  | <u>519</u> | <u>35</u> | <u>16</u> |  |
| <u>2-OHE2-bisDS</u> | <u>521.2</u> | <u>755.2</u> | <u>81</u> | <u>10</u> | <u>521</u> | <u>37</u> | <u>16</u> | <u>17.93</u> |
|  |  |  |  |  | <u>170</u> | <u>47</u> | <u>8</u> |  |
| <u>4-OHE1-bisDS</u> | <u>519.19</u> | <u>753.1</u> | <u>81</u> | <u>10</u> | <u>707</u> | <u>11</u> | <u>6</u> | <u>18.06</u> |
|  |  |  |  |  | <u>685</u> | <u>21</u> | <u>28</u> |  |
| <u>16α-OHE1-DS</u> | <u>519.19</u> | <u>520.2</u> | <u>101</u> | <u>10</u> | <u>115</u> | <u>109</u> | <u>16</u> | <u>15.32</u> |
|  |  |  |  |  | <u>156</u> | <u>55</u> | <u>8</u> |  |
| <u>3-MeOE1-DS</u> | <u>533.2</u> | <u>534.2</u> | <u>81</u> | <u>10</u> | <u>171</u> | <u>41</u> | <u>8</u> | <u>16.69</u> |
|  |  |  |  |  | <u>156</u> | <u>83</u> | <u>20</u> |  |
| <u>4-MeOE1-DS</u> | <u>533.2</u> | <u>534.2</u> | <u>101</u> | <u>10</u> | <u>171</u> | <u>41</u> | <u>8</u> | <u>17.07</u> |
|  |  |  |  |  | <u>156</u> | <u>77</u> | <u>8</u> |  |
| <u>17-epiE3-DS</u> | <u>521.2</u> | <u>522.2</u> | <u>91</u> | <u>10</u> | <u>171</u> | <u>47</u> | <u>8</u> | <u>15.6</u> |
|  |  |  |  |  | <u>156</u> | <u>57</u> | <u>8</u> |  |
| <u>2-MeOE1-DS</u> | <u>533.2</u> | <u>534.2</u> | <u>71</u> | <u>10</u> | <u>171</u> | <u>41</u> | <u>8</u> | <u>16.9</u> |
|  |  |  |  |  | <u>156</u> | <u>83</u> | <u>18</u> |  |
| <u>2-MeOE2-DS</u> | <u>535.22</u> | <u>536.2</u> | <u>116</u> | <u>10</u> | <u>171</u> | <u>43</u> | <u>8</u> | <u>16.58</u> |
|  |  |  |  |  | <u>156</u> | <u>79</u> | <u>24</u> |  |
| <u>4-MeOE2-DS</u> | <u>535.22</u> | <u>536.2</u> | <u>96</u> | <u>10</u> | <u>171</u> | <u>41</u> | <u>24</u> | <u>16.71</u> |
|  |  |  |  |  | <u>156</u> | <u>81</u> | <u>20</u> |  |
| <u>16-epiE3-DS</u> | <u>521.2</u> | <u>522.2</u> | <u>76</u> | <u>10</u> | <u>171</u> | <u>45</u> | <u>8</u> | <u>15.36</u> |
|  |  |  |  |  | <u>156</u> | <u>55</u> | <u>22</u> |  |
| <u>16-ketoE2-DS</u> | <u>519.19</u> | <u>520.2</u> | <u>76</u> | <u>10</u> | <u>171</u> | <u>43</u> | <u>8</u> | <u>15.21</u> |
|  |  |  |  |  | <u>156</u> | <u>55</u> | <u>8</u> |  |
| <u>E2-D4-DS</u> | <u>509.21</u> | <u>510.3</u> | <u>51</u> | <u>10</u> | <u>171</u> | <u>47</u> | <u>10</u> | <u>16.77</u> |
|  |  |  |  |  | <u>156</u> | <u>57</u> | <u>20</u> |  |
| <u>E1-D4-DS</u> | <u>507.19</u> | <u>508.2</u> | <u>96</u> | <u>10</u> | <u>171</u> | <u>43</u> | <u>8</u> | <u>17.06</u> |
|  |  |  |  |  | <u>156</u> | <u>57</u> | <u>18</u> |  |

| <u>Analyte</u> | <u>Mmi</u> | <u>Q1 mass (Da)</u> | <u>DP (v)</u> | <u>EP (v)</u> | <u>Q3 mass (Da)</u> | <u>CE (v)</u> | <u>CXP (v)</u> | <u>RT (min)</u> |
| --- | --- | --- | --- | --- | --- | --- | --- | --- |
| <u>E1-DS-D6</u> | <u>509.19</u> | <u>510.2</u> | <u>80</u> | <u>10</u> | <u>177</u> | <u>45</u> | <u>10</u> | <u>16.99</u> |
|  |  |  |  |  | <u>159</u> | <u>59</u> | <u>18</u> |  |
| <u>E2-DS-D6</u> | <u>511.21</u> | <u>512.2</u> | <u>136</u> | <u>10</u> | <u>177</u> | <u>47</u> | <u>10</u> | <u>16.72</u> |
|  |  |  |  |  | <u>159</u> | <u>59</u> | <u>18</u> |  |
| <u>E3-DS-D6</u> | <u>527.2</u> | <u>528.2</u> | <u>71</u> | <u>10</u> | <u>177</u> | <u>47</u> | <u>8</u> | <u>14.15</u> |
|  |  |  |  |  | <u>159</u> | <u>57</u> | <u>4</u> |  |
| <u>2-OHE1-bisDS-D6</u> | <u>764.19</u> | <u>765.2</u> | <u>61</u> | <u>10</u> | <u>176</u> | <u>45</u> | <u>10</u> | <u>18.02</u> |
|  |  |  |  |  | <u>525</u> | <u>37</u> | <u>18</u> |  |
| <u>2-OHE2-bisDS-D6</u> | <u>766.2</u> | <u>767.2</u> | <u>96</u> | <u>10</u> | <u>176</u> | <u>49</u> | <u>8</u> | <u>17.83</u> |
|  |  |  |  |  | <u>527</u> | <u>39</u> | <u>8</u> |  |
| <u>4-OHE1-bisDS-D6</u> | <u>764.19</u> | <u>765.2</u> | <u>61</u> | <u>10</u> | <u>176</u> | <u>45</u> | <u>10</u> | <u>17.96</u> |
|  |  |  |  |  | <u>525</u> | <u>37</u> | <u>18</u> |  |
| <u>16α-OHE1-DS-D6</u> | <u>525.19</u> | <u>526.2</u> | <u>86</u> | <u>10</u> | <u>177</u> | <u>45</u> | <u>8</u> | <u>15.22</u> |
|  |  |  |  |  | <u>159</u> | <u>57</u> | <u>8</u> |  |
| <u>3-MeOE1-DS-D6</u> | <u>539.2</u> | <u>540.2</u> | <u>66</u> | <u>10</u> | <u>177</u> | <u>43</u> | <u>8</u> | <u>16.56</u> |
|  |  |  |  |  | <u>159</u> | <u>85</u> | <u>8</u> |  |
| <u>4-MeOE1-DS-D6</u> | <u>539.2</u> | <u>540.2</u> | <u>91</u> | <u>10</u> | <u>177</u> | <u>41</u> | <u>8</u> | <u>16.97</u> |
|  |  |  |  |  | <u>159</u> | <u>83</u> | <u>20</u> |  |
| <u>17-epiE3-DS-D6</u> | <u>527.2</u> | <u>528.2</u> | <u>176</u> | <u>10</u> | <u>177</u> | <u>47</u> | <u>8</u> | <u>15.48</u> |
|  |  |  |  |  | <u>159</u> | <u>57</u> | <u>8</u> |  |
| <u>2-MeOE1-DS-D6</u> | <u>539.2</u> | <u>540.2</u> | <u>106</u> | <u>10</u> | <u>177</u> | <u>43</u> | <u>8</u> | <u>16.79</u> |
|  |  |  |  |  | <u>159</u> | <u>75</u> | <u>10</u> |  |
| <u>2-MeOE2-DS-D6</u> | <u>541.2</u> | <u>542.2</u> | <u>106</u> | <u>10</u> | <u>177</u> | <u>45</u> | <u>8</u> | <u>16.46</u> |
|  |  |  |  |  | <u>159</u> | <u>75</u> | <u>10</u> |  |
| <u>4-MeOE2-DS-D6</u> | <u>541.2</u> | <u>542.2</u> | <u>51</u> | <u>10</u> | <u>177</u> | <u>43</u> | <u>8</u> | <u>16.61</u> |
|  |  |  |  |  | <u>159</u> | <u>75</u> | <u>10</u> |  |
| <u>16-epiE3-DS-D6</u> | <u>527.2</u> | <u>528.2</u> | <u>71</u> | <u>10</u> | <u>177</u> | <u>47</u> | <u>8</u> | <u>15.25</u> |
|  |  |  |  |  | <u>159</u> | <u>61</u> | <u>8</u> |  |
| <u>16-ketoE2-DS-D6</u> | <u>525.19</u> | <u>526.2</u> | <u>96</u> | <u>10</u> | <u>177</u> | <u>45</u> | <u>8</u> | <u>15.11</u> |
|  |  |  |  |  | <u>159</u> | <u>57</u> | <u>8</u> |  |

